# Genetic risk for major depressive disorder and loneliness in gender-specific associations with coronary artery disease

**DOI:** 10.1101/512541

**Authors:** Jessica Dennis, Julia Sealock, Rebecca T Levinson, Eric Farber-Eger, Jacob Franco, Sarah Fong, Peter Straub, Donald Hucks, MacRae F Linton, Wen-Liang Song, Pierre Fontanillas, Sarah L Elson, Douglas Ruderfer, Abdel Abdellaoui, Sandra Sanchez-Roige, Abraham A Palmer, Dorret I Boomsma, Nancy J Cox, Guanhua Chen, Jonathan D Mosley, Quinn S Wells, Lea K Davis

**Affiliations:** Division of Genetic Medicine, Department of Medicine Vanderbilt University Medical Center Nashville, TN; Vanderbilt Genetics Institute Vanderbilt University Medical Center Nashville, TN; Division of Cardiovascular Medicine, Department of Medicine Vanderbilt University Medical Center Nashville, TN; Vanderbilt Institute for Clinical and Translational Research Vanderbilt University Medical Center Nashville, TN; Division of General Internal Medicine and Public Health, Department of Medicine Vanderbilt University Medical Center Nashville, TN; 23andMe, Inc. Mountain View, CA; Department of Biological Psychology Vrije Universiteit Amsterdam, The Netherlands; Department Psychiatry, Amsterdam UMC, University of Amsterdam Amsterdam, The Netherlands; Department of Psychiatry University of California San Diego La Jolla, CA; Institute for Genomic Medicine University of California San Diego La Jolla, CA; Netherlands Twin Register, Department of Biological Psychology Vrije Universiteit Amsterdam, The Netherlands; APH (Amsterdam Public Health) Institute Amsterdam, The Netherlands; Department of Biostatistics and Medical lnformatics University of Wisconsin-Madison Madison, WI; Departments of Medicine and Biomedical Informatics Vanderbilt University Medical Center Nashville, TN

**Author notes:** Corresponding Author: Lea K Davis, PhD, Assistant Professor of Medicine, Vanderbilt Genetics Institute, Division of Genetic Medicine, Department of Medicine, 511-A Light Hall, Vanderbilt University, 2215 Garland Ave, Nashville TN 37232.

## Abstract

**Importance:** Epidemiological evidence indicates that major depressive disorder (MDD) and loneliness both reduce life expectancies, but mechanisms underlying the excess morbidity are unclear. Electronic health records (EHRs) linked to genetic data offer new opportunities to address this knowledge gap.

**Objective:** To determine the medical morbidity pattern associated with genetic risk factors for MDD and loneliness, two common psychological traits with adverse health outcomes.

**Design:** Phenome-wide association study using EHRs spanning 1990 to 2017 from the Vanderbilt University Medical Center biobank, BioVU. Top associations with coronary artery disease (CAD) were replicated in the Atherosclerosis Risk in Communities (ARIC) cohort.

**Setting:** Hospital-based EHR study, with replication in a population-based cohort study.

**Participants:** 18,385 genotyped adult patients in BioVU. Replication in ARIC included 7,197 genotyped participants. All participants were of European ancestry.

**Exposures:** Polygenic scores for MDD and loneliness were developed for each individual using previously published meta-GWAS summary statistics.

**Main Outcomes and Measures:** The phenome-wide association study included 882 clinical diagnoses ascertained via billing codes in the EHR. ARIC included 1598 incident CAD cases.

**Results:** BioVU patients had a median EHR length of 9.91 years. In the phenome-wide association study, polygenic scores for MDD and loneliness were significantly associated with psychiatric and cardiac phenotypes. Targeted analyses of CAD in 3,893 cases and 4,197 controls in BioVU found odds ratios of 1.11 (95% CI, 1.04-1.18; P=8.43×10^−4^) and 1.13 (95% CI, 1.07-1.20; P=4.51×10^−6^) per 1-SD increase in the polygenic scores for MDD and loneliness, respectively. Comparable hazard ratios in ARIC were 1.07 (95% CI, 0.99-1.14; P=0.07) and 1.07 (1.01-1.15; P=0.03). Across both studies, the increased risk persisted in women after adjusting for multiple conventional risk factors, a polygenic score for CAD, and psychiatric symptoms (available in BioVU). Controlling for genetic risk factors shared between MDD and loneliness, the polygenic score for loneliness conditioned on MDD remained associated with CAD risk, but the polygenic score for MDD conditioned on loneliness did not.

**Conclusions and Relevance:** Genetic risk factors for MDD and loneliness act pleiotropically to increase CAD risk in women. Continued research into the biological and clinical connections between the heart and mind is warranted.

## Introduction

Major depressive disorder (MDD) has a lifetime prevalence of 17%^1^ and is associated with a 71% increased risk of premature death.^2^ Suicide and injury account for some of the excess mortality, yet most people with MDD die of common, chronic disease, including heart disease.^2,3^ Recent evidence also implicates chronic loneliness, i.e., the discrepancy between a person’s desired and perceived social connectedness, in early mortality.^4–6^ Chronic loneliness is experienced by up to 22% of adults^7^ and can lead to MDD, however, the two traits are only moderately phenotypically correlated and are believed to represent distinct underlying phenomena.^8,9^

Disease risk factors such as obesity and smoking are more common in people who are depressed and lonely,^3,4^ but few studies have examined the proportion of early mortality risk explained by these risk factors,^10^ or whether risks differ by sex.^10–13^ Electronic health records (EHRs) contain dense phenotype data and provide an opportunity to comprehensively study disease risk factors and comorbidity patterns across patient groups. Additionally, when linked to genotype information, EHRs facilitate use of genetic methods such as polygenic scoring. Polygenic scores model the continuous genetic liability to traits, and can be calculated in any individual with genetic data, even if the trait itself is not measured.

This study included five aims. The first was to identify the medical morbidity patterns associated with polygenic scores for MDD and loneliness in the Vanderbilt University Medical Center (VUMC) EHR and accompanying biobank, BioVU. Heart disease and depression were the diagnoses most strongly associated with both polygenic scores, providing rationale for the four remaining aims of the study. Second, we undertook a targeted analysis of coronary artery disease (CAD), the most common form of heart disease, and determined whether the observed associations were a consequence of depression, or the result of pleiotropic genetic variants. Third, we examined whether the increased risk of CAD could be attenuated by accounting for multiple conventional CAD risk factors. Fourth, given the reported sex differences in mortality, presentation, and risk factors,^14,15^ we repeated all analyses in males and females separately. Fifth, we replicated our findings in a large prospectively collected population cohort, the Atherosclerosis Risk in Communities (ARIC) study. Our results provide strong evidence that pleiotropic risk factors for MDD and loneliness increase the risk of CAD, with more pronounced effects in females than in males.

## Methods

### Study Populations

This project was approved by the VUMC Institutional Review Board (IRB #160302). The VUMC biobank, BioVU, served as the discovery population. BioVU includes more than 285,000 patients whose DNA was extracted from discarded blood, and linked to de-identified EHR data spanning 1990 to 2017.^16^ The Atherosclerosis Risk in Communities (ARIC) cohort served as our replication population.^17^ The study included 15,792 participants aged 45 to 64 years in 1987 recruited from four geographic areas in the US, assessed at baseline for heart disease risk factors, and followed until December 31, 2004 for health outcomes.^18^

### Genotyping and Quality Control

A subset of BioVU patients (N=24,262) was genotyped as part of various institutional and investigator-initiated projects on the Illumina MEGA^EX^ platform, which contains more than 2 million markers. Quality control proceeded as previously described.^19^ Genotypes were imputed using SHAPEIT^20^/IMPUTE4^21^ with the 1000 genomes phase I reference panel, and variants with INFO <.3 were excluded. A subset of SNPs in linkage disequilibrium was used to calculate relatedness and principal components of ancestry using multidimensional scaling in PLINK v1.9.^22^ We randomly removed one individual from pairs of highly related individuals (pihat >.1) to avoid spurious results driven by cryptic relatedness, and restricted to a homogenous population of European descent defined by principal components of ancestry to avoid population stratification effects, leaving 18,385 individuals for analyses. Samples were genotyped in five batches, and variants were removed if allele frequencies differed significantly (P < 5×10^−5^) between any batch and the rest of the sample. Finally, we filtered multi-allelic and structural variants, converted dosage data to hard genotype calls, and excluded variants with uncertainty >.1 or INFO <.95, resulting in 5,218,407 high quality SNPs across the autosomes.

Genetic data from 13,113 ARIC participants were downloaded from dbGaP (phs000280.v3.p1). Genotypes were measured on the Affymetrix 6.0 SNP array, and data were cleaned as previously described.^23^ Genetic ancestry was assigned using STRUCTURE^24^ in conjunction with HapMap reference populations, and European ancestry was defined as >90% probability of being in the CEU cluster. Genotypes were imputed to the October 2014 Phase 3 release of the 1000 Genomes cosmopolitan reference haplotypes^25^ using SHAPEIT^20^/IMPUTE2,^21^ and we excluded SNPs with INFO <.7, multi-allelic SNPs, and structural variants. Dosage data were converted to hard genotyping calls, filtering variants with uncertainty >.1, leaving 6,569,625 high quality SNPs for analysis. Inter-individual relatedness and principal components of ancestry were calculated by GCTA,^26^ and we randomly excluded one individual from pairs of related individuals (pihat >.05), leaving 6,975 unrelated European ancestry participants.

### Phenotype Data

To facilitate investigation of comorbidity patterns across the medical phenome, we mapped International Classification of Diseases, 9^th^ edition (ICD-9) billing codes in the EHR to phecodes, which are higher-order representations of disease categories. ICD-9 codes were mapped to 1814 phecode categories according to the Phecode Map v1.2 (https://phewascatalog.org/phecodes), as implemented in the PheWAS R package v0.12.^27^ Phecodes required two or more occurrences of a component ICD-9 code assigned on different days,^28^ a validated method to improve the positive predictive value of phecodes,^28^ and the control group excluded patients with only one component ICD-9 code, or with one or more ICD-9 codes that mapped to related phecodes (as defined by the Phecode Map).

Heart disease and its most common manifestation, CAD, was a focus of our analysis. However, no single phecode was diagnostic of CAD. Therefore, we defined CAD in BioVU patients by a random forest (machine learning) classifier^29^ that integrated data from across the EHR (eMethods; eFigure 1; eFigure 2; eFigure 3). Age in cases was defined by the age of first CAD-defining feature in the EHR, while in controls, age was defined by the age at last encounter. Data on CAD risk factors were also extracted from the EHR either from structured data or via text-mining algorithms (eMethods). CAD risk factors included body mass index (BMI), hypertension, type 2 diabetes diagnosis, pre-medication blood levels of high- and low-density lipoprotein cholesterol (HDL and LDL) and of triglycerides^30^ (lipid-altering medications are listed in eTable 1), smoking history, and socio-economic status.

A key research question was whether associations with polygenic scores for MDD and loneliness persisted in people without a clinical MDD diagnosis. We defined MDD by the phecode 296.22 “Major depressive disorder”, which included both single episode and recurrent major depressive episodes that are severe, moderate, or of unspecified severity. To capture patients with depressive symptoms that may not meet criteria for MDD, or that may have lacked an MDD diagnosis in their EHR, we also defined a milder depressive symptoms phenotype by one or more depression ICD-9 codes (296.2 and 296.3, including all fifth digit subclassifications, and 311). As a final test of the robustness of associations, we excluded patients with any psychiatric symptoms (eMethods).

Phenotype data for ARIC samples came from measurements aggregated by the GENEVA substudy (pht000114.v2.p1), downloaded with permission from dbGaP.^23^ Eligible participants were those with data on incident CAD, defined as myocardial infarction, fatal coronary heart disease, silent myocardial infarction detected by electrocardiography, or revascularization procedure. We additionally extracted data on sex, age at first visit, and time to event (CAD or censoring in controls), as well as first visit data on BMI, waist girth, smoking status, hypertension medication use, systolic and diastolic blood pressure, type 2 diabetes diagnosis, highest level of education, use of cholesterol-lowering medication and other medications that secondarily affect cholesterol, and blood measurements of HDL, LDL, and triglycerides (eTable 2). We defined hypertension by the variable hypertension medication use, or systolic blood pressure > 130 mmHg, or diastolic blood pressure > 80 mmHg.^31^ Depression was assessed in 2011-2013 via the centers for epidemiological studies-depression scale, but these data exist in only 6,471 of the original 15,792 ARIC cohort members^18^ and so were not included in this analysis.

### Statistical Analyses

Polygenic scores for MDD and loneliness were computed using the PRSice software package^32^ (eMethods) for each individual in BioVU and ARIC as follows:

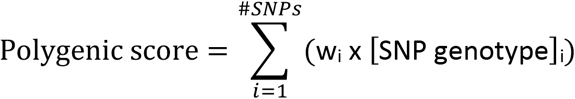

where the SNP genotype was coded as 0, 1 or 2 and w_i_ was the ß coefficient (representing effect size of the marginal association between the SNP and the phenotype) from the Psychiatric Genomics Consortium meta-GWAS of MDD,^33^ or the Lonely Consortium meta-GWAS of loneliness.^34^ We selected SNPs into the polygenic score if their association *P* in the meta-GWAS was below a specified threshold. In the phenome-wide association study, we selected an *P* threshold of.05, a value previously found to generate scores that maximized the prediction of MDD^33^ and other psychiatric traits,^35^ and included 392,372 and 379,906 SNPs in the polygenic scores of MDD and loneliness, respectively. In targeted analyses of CAD, we selected the *P* threshold at which the polygenic score was most strongly associated with CAD risk. This approach improved our power to detect associations with CAD by maximizing the number of pleiotropic SNPs included in the MDD and loneliness polygenic scores. We determined *P* thresholds of 9.0005×10^−4^ for MDD (n SNPs=1,387) and 2.50005×10^−3^ for loneliness (n SNPs=2,253) by iterating over *P* from 5×10^−8^ to 1 in increments of 5×10^−5^ and assessing fit via Nagelkerke’s pseudo R^2^ (eFigure 4). Polygenic scores calculated at these best fit *P* thresholds were used in all sensitivity and replication analyses of CAD to minimize false findings due to over-fitting. To adjust for the genetic correlation between MDD and loneliness (r_g_=.64; SE=.03; P=2.8×10^−114^),^34^ we used the multi-trait-based conditional and joint analysis (mtCOJO) package,^36^ which uses only meta-GWAS summary statistics to statistically isolate independent genetic associations. Using mtCOJO, we calculated SNP associations with MDD conditional on loneliness (MDD | loneliness), and vice-versa (loneliness |MDD), and calculated polygenic scores from the MDD |loneliness and loneliness |MDD summary statistics using the best fit *P* thresholds above. All polygenic scores were generated by the default pruning and thresholding parameters in PRSice v2^32^ (eMethods) and were scaled to have an SD of 1. Risk ratios (odds or hazard) reflect the risk increase per 1-SD increase in the polygenic score.

Phenome-wide association studies were conducted with the PheWAS R package v0.12.^27^ We required phecodes to have at least 100 cases, and included covariates for sex, median age across the EHR, genotyping batch, and the first 10 principal components of ancestry.

Targeted analyses of CAD in BioVU employed logistic regression. Minimally adjusted models included covariates for sex, age, genotype batch, and the first 10 principal components of ancestry. Fully adjusted models additionally included covariates for BMI, hypertension, smoking, type 2 diabetes, blood measurements of HDL, LDL, and triglycerides, highest level of education, a polygenic score for CAD (eMethods), and depressive symptoms. In ARIC, we modeled incident CAD using a Cox proportional hazards model and included sex, baseline age, and the first 10 principal components of ancestry in the minimally adjusted models. Fully adjusted models included additional covariates for waist girth, smoking status, hypertension medication use, systolic blood pressure, type 2 diabetes, highest level of education, use of cholesterol-lowering medication and other medications that secondarily affect cholesterol, and blood measurements of HDL, LDL, and triglycerides. Although these covariates did not exactly match those we controlled for in BioVU, (i.e., waist girth instead of BMI, hypertension medication use and systolic blood pressure instead of hypertension diagnosis), we used them preferentially since they are better measures of the underlying risk factors.

All analyses were completed in R version 3.4.3 or 3.5.

## Results

Our analyses included 18,385 BioVU patients. These patients had a median EHR length of 9.91 years (range less than one year to 27 years; Table 1), the average median age across the EHR was 57.2 years (SD 17.6), and 50.9% of patients were female. Nearly 5% of patients were diagnosed with MDD, 24.9% had depressive symptoms, and 39.5% had evidence of any psychiatric symptoms.

**Table 1.** Characteristics of genotyped BioVU patients included in the phenome-wide association study.

|  | **BioVU (N=18,385)** |
| --- | --- |
| Length of medical record in years, median (min, max) | 9.91 (0, 27.06) |
| Median age across medical record, mean (sd) | 57.20 (17.58) |
| Female, No. (%) | 9355 (50.9) |
| MDD diagnosis, No. (%) | 826 (4.5) |
| Depressive symptoms, No. (%) | 4576 (24.9) |
| Any psychiatric symptoms, No. (%) | 7267 (39.5) |

MDD denotes major depressive disorder.

In phenome-wide association studies, we analyzed 882 phecodes with at least 100 cases, and used a Bonferroni-corrected phenome-wide significance threshold of.05/882=5.67×10^−5^. This threshold, however, is likely over-conservative because it incorrectly assumes independence between phecodes. The phenome-wide association study of the polygenic score for MDD identified phenome-wide significant associations with “Mood disorders” (odds ratio [OR], 1.10 [95% CI, 1.06-1.15]; P=1.83×10^−6^), and “Depression” (OR, 1.10 [95% CI, 1.05-1.15]; P=1.13×10^−5^), parent codes for MDD, as expected (Figure 1A; interactive plots available at: https://sealockj.shinyapps.io/mdd_loneliness_interactive/). The remaining six phecodes that were significant after Bonferroni correction were related to acute and chronic heart diseases and their associated risk factors, including “Ischemic heart disease” (OR, 1.09 [95% CI, 1.05-1.14]; P=8.28×10-6) and “Coronary atherosclerosis” (OR, 1.10 [95% CI, 1.05-1.14]; P=1.08×10-5). The phenome-wide association study of the polygenic score for loneliness (Figure 1B) also found significant associations with “Mood disorders” (OR, 1.09 [95% CI, 1.05-1.14]; P=1.39×10^−5^) and “Depression” (OR, 1.10 [95% CI, 1.05-1.14]; P=1.43×10^−5^). The top associations, however, were with heart disease phenotypes, specifically, “Ischemic heart disease” (OR, 1.10 [95% CI, 1. 06-1.14]; P=2.91×10^−6^) and “Coronary atherosclerosis” (OR, 1.10 [95% CI, 1.06-1.15]; P=5.89×10^−6^).

**Figure 1.**
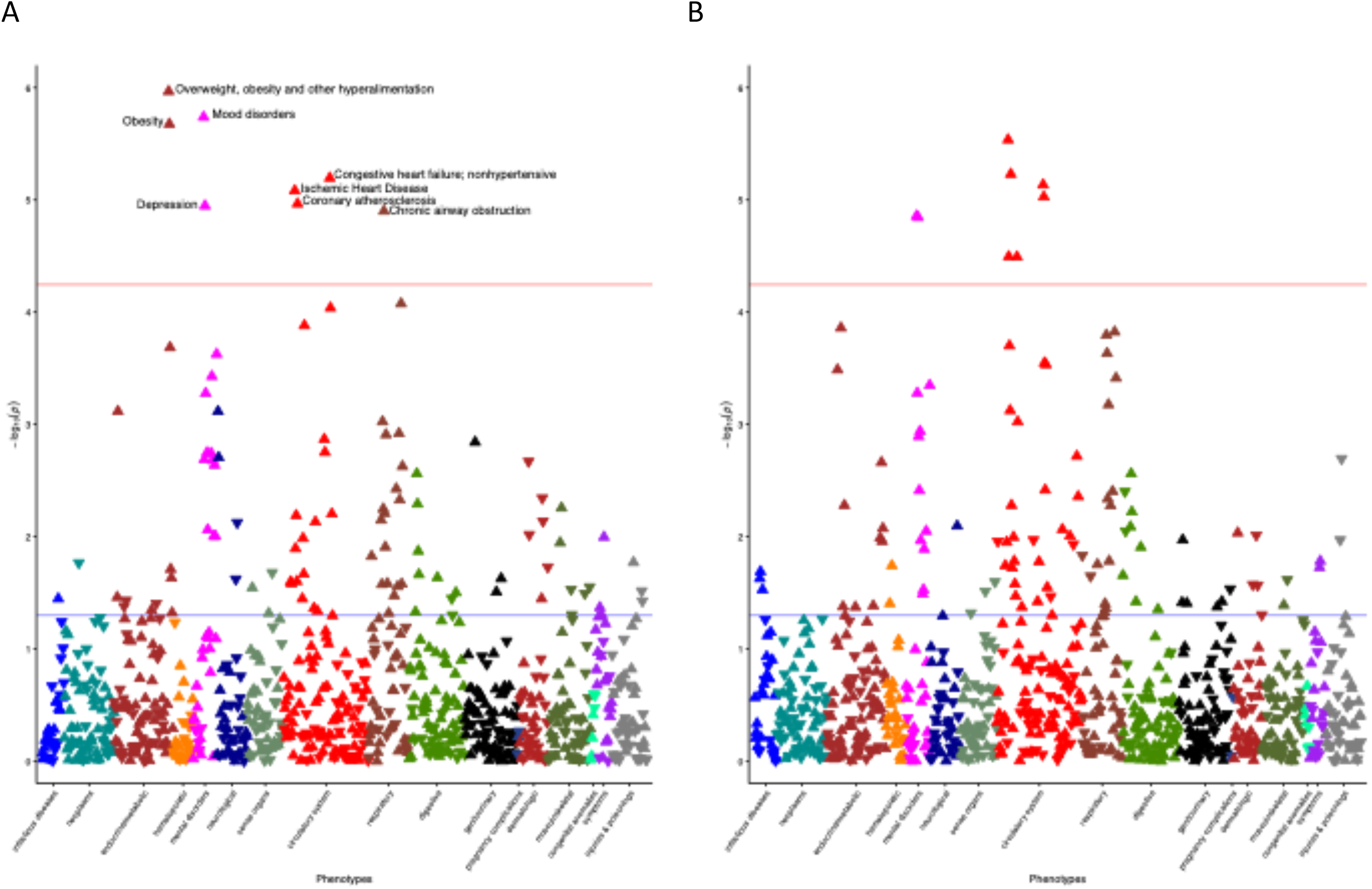
Results from phenome-wide association studies of polygenic scores for MDD (A) and loneliness (B) in BioVU. The red line denotes the Bonferroni threshold for statistical significance (.05/882=5.67×10^−5^), and phenome-wide significant phecodes are labelled. Upward triangles indicate increased odds for a given phecode per 1-SD increased risk in the polygenic score, while downward triangles indicate reduced odds of a given phecode.

Motivated by these findings, we targeted CAD, the most common manifestation of heart disease, to better understand the contribution of genetic risk factors for MDD and loneliness to their shared comorbidity patterns implicating CAD. Our machine learning algorithm identified 3,893 cases and 4,197 controls in BioVU (eTable 3). In minimally adjusted models, each SD increase in the polygenic scores for MDD and loneliness respectively increased the odds of CAD by 1.11 (95% CI, 1.04-1.18; P=8.43×10^−4^) and 1.13 (95% CI, 1.08-1.20; P=4.51×10^−6^). Stratified by deciles, patients with polygenic scores for MDD and loneliness in the top versus bottom deciles had, respectively, a 1.53-fold (95% CI, 1.18-1.98; P=1.2×10^−3^) and 1.51-fold (95%CI, 1.19-1.91; P=7.4×10^−4^) greater risk of CAD (Figure 2). These associations persisted even after excluding patients with a clinical diagnosis of MDD, depressive symptoms, or any psychiatric symptoms (Table 2).

**Figure 2.**
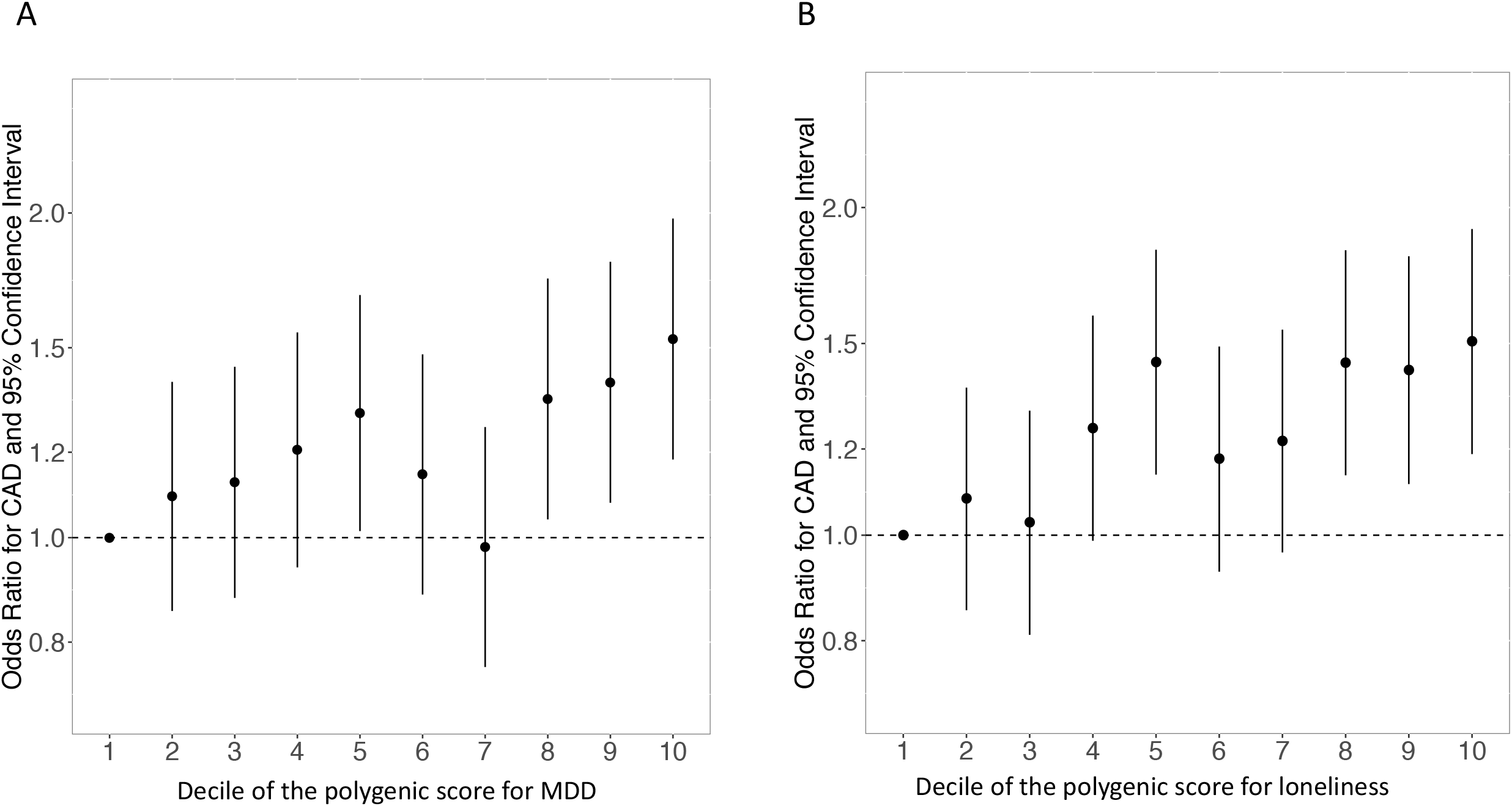
Odds of CAD by decile of the polygenic score for MDD (A) and loneliness (B) in BioVU. The referent group in all calculations is the lowest polygenic score decile.

**Table 2.** Associations between polygenic scores for major depressive disorder and loneliness with coronary artery disease after excluding patients with a diagnosis of major depressive disorder, depressive symptoms, or any psychiatric symptoms.

| Polygenic score | Patient exclusions | N Cases | N Controls | OR (95% CI) | P |
| --- | --- | --- | --- | --- | --- |
| MDD | MDD | 3744 | 4037 | 1.11 (1.05-1.18) | $6.54 \times 10^{-4}$ |
| MDD | Depressive symptoms | 2935 | 3147 | 1.10 (1.03-1.18) | $5.76 \times 10^{-3}$ |
| MDD | Any psychiatric symptoms | 2440 | 2551 | 1.11 (1.03-1.19) | $9.27 \times 10^{-3}$ |
| Loneliness | MDD | 3744 | 4037 | 1.13 (1.07-1.20) | $9.22 \times 10^{-6}$ |
| Loneliness | Depressive symptoms | 2935 | 3147 | 1.14 (1.08-1.22) | $2.08 \times 10^{-5}$ |
| Loneliness | Any psychiatric symptoms | 2440 | 2551 | 1.12 (1.05-1.20) | $8.69 \times 10^{-4}$ |

MDD denotes major depressive disorder

Associations were also robust to adjustment for conventional risk factors in women, but not in men (Figures 3A and 3B). We next accounted for the genetic correlation between MDD and loneliness using the mtCOJO package.^36^ The polygenic score for MDD |loneliness was not associated with CAD risk (Figure 3C), whereas the score for loneliness |MDD remained associated with CAD risk in women, even in the fully adjusted model (Figure 3D). The polygenic score for loneliness |MDD was associated with a 1.23-fold (95% CI, 1.02-1.47; P=.017) increased odds of CAD in women in the fully adjusted model that maximized patient inclusion by including categories for missing values. The association was equivalent, although less statistically significant (OR, 1.23 [95% CI, 0.94-1.60]; P=.126) in the fully adjusted model that excluded the 699 female BioVU patients with missing covariate data.

**Figure 3.**
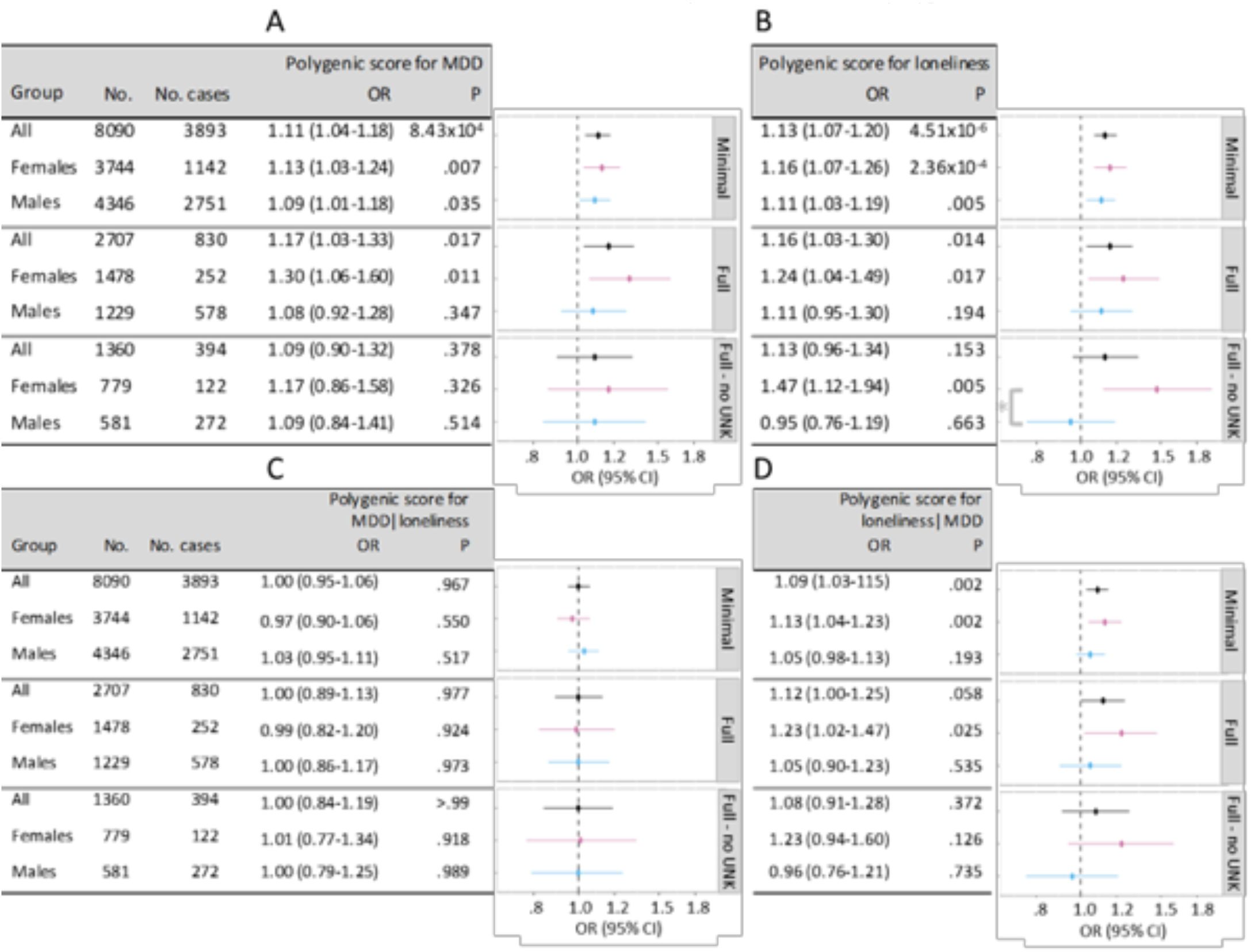
Risk of CAD predicted by polygenic scores for MDD (A), loneliness (B), MDD | loneliness (C), and loneliness |MDD (D) before and after adjustment for conventional heart disease risk factors. Minimally adjusted (“Minimal”) models included sex (except sex-stratified results), age, the first 10 principal components of ancestry, and genotype batch. Fully adjusted (“Full”) models included additional covariates for BMI, hypertension, smoking, type 2 diabetes, blood measurements of HDL, LDL, and triglycerides, highest level of education, a polygenic score for CAD, and depressive symptoms. Values for smoking and highest level of education were unknown for ~50% of patients. These unknown values were retained in the “Full” model by modeling an explicit category for “missing” values, and were removed from the “Full - no UNK” model. The asterisk in panel D denotes sex*polygenic score interaction P <.05.

The ARIC sample included 7,197 participants of European ancestry with clean genotyping data, followed for a median of 16.0 years (range 0.05 to 18.1), over which time 923 incident CAD cases were recorded (eTable 4). The polygenic score for MDD was associated with a hazard ratio (HR) of 1.07 (95% CI, 0.99-1.14; P=.07), as was the polygenic score for loneliness (HR, 1.07 [95% CI, 1.01-1.15]; P=.03). Associations were robust to confounder adjustment in women but not in men (eFigure 5), and the polygenic score for loneliness |MDD was associated with a CAD HR of 1.14 (95% CI, 1.01-1.29; P=.02) in women in the fully adjusted model.

## Discussion

This study investigated the phenome-wide consequences of genetic risk factors for MDD and loneliness in BioVU, which is a deeply phenotyped large-scale EHR collection with linked genotype data. We discovered strong associations between polygenic scores for MDD and loneliness with heart disease, and our targeted analysis of CAD revealed that patients with polygenic scores in the highest decile for either MDD or loneliness had a more than 50% greater risk of CAD compared to patients with scores in the lowest decile. An epidemiological association between MDD and heart disease is known;^37^ here we showed that the association is partly attributable to genetics. Similar results were observed in ARIC participants, who were prospectively surveyed for heart disease outcomes using traditional epidemiological approaches.

An unique feature of our study was the extensive sensitivity analyses that provided more insight into the reported genetic and epidemiological correlations between MDD, loneliness, and CAD. Genetic correlations alone can be difficult to interpret because they may be influenced by “phenotypic hitchhiking”. For example, if MDD is comorbid with heart disease, then a GWAS of MDD will also ascertain cases with heart disease, which could then induce a genetic correlation between MDD and an independent GWAS of heart disease. In this study, we controlled for multiple clinical risk factors and known CAD risk variants, resulting in strong evidence that MDD and loneliness risk variants exert pleiotropic effects on CAD.

MDD and loneliness are genetically correlated.^34^ Our conditional analyses, however, suggested that genetic risk factors specific to loneliness increased CAD risk independent of the genetic risk factors that are shared between MDD and loneliness. This finding is congruent with the physiological effects of chronic loneliness in humans and in animal models.^38^ Loneliness induces a state of self-preservation in anticipation of being without the protection of others: it triggers depressive symptoms that signal the need for support and connection from peers;^9^ disrupts sleep to maintain a state of alertness at night;^39^ raises blood pressure;^40^ and activates the hypothalamic pituitary adrenal axis,^38,41–43^ which regulates cortisol, a key hormone in stress reactions, metabolism, digestion, immunity, and energy storage. While these responses may be advantageous for being alone in the short term, health problems ensue when loneliness is chronic and the biological response is sustained.^38^

Both MDD and heart disease exhibit significant sex-differences in prevalence and presentation,^1,14^ and heart disease remains the number one killer of women in the United States.^15^ Our analyses revealed that genetic risk for MDD and loneliness conferred a higher risk of CAD in women than in men, and that the risk in women was robust to adjustment for the major known risk factors. There are two possible, non-mutually exclusive, explanations for this finding. First, genetic predisposition to MDD or loneliness may be a chronic risk factor for females but an acute risk factor for males, as was previously suggested in a study of 30,000 twins from the Swedish population-based twin registry.^11^ Second, the conventional risk factors identified in most epidemiological studies may be poorer predictors of CAD for females than they are for males. Future work is warranted to investigate the role of pleiotropic genes on both heart disease and MDD, especially in females where the effects are strongest, and the etiology of heart disease more obscure.

### Limitations

EHR data reflect real clinical use patterns, but some data can be missing or inaccurate. We addressed this potential limitation by implementing a carefully curated CAD phenotype, and conducted sensitivity analyses excluding people with unknown covariate values. We also replicated our findings in a prospective epidemiological cohort study. A second limitation of our study was its restriction to individuals of European ancestry, but this decision was necessitated by the ancestry of patients in the meta-GWAS used to build the polygenic scores. The relevance of our findings to individuals of other ancestries, therefore, is unknown.^44,45^

## Conclusions

Mental health has historically been siloed away from the rest of medicine, resulting in a poor understanding of the relationship between mental and physical health. This EHR-based study showed that genetic risk for MDD and loneliness were just as strongly associated with heart disease as with depression itself. Moreover, the increased risk of CAD persisted in women after adjusting for psychiatric symptoms and multiple other risk factors, suggesting that the excess risk is due to pleiotropic genetic effects and is not simply a behavioral consequence of depression. Identification of these pleiotropic genes could advance both precision medicine and drug development.

## Acknowledgements

### Author Contributions

JD, QSW, and LKD conceived of and designed the study. All authors contributed to the acquisition, analysis, or interpretation of data. JD, JDM, QSW, and LKD drafted the manuscript. All authors contributed to critical revision of the manuscript for intellectual content.

### Funding/Support

JD is supported by the Canadian Institutes of Health Research (award MFE-142936). RTL is supported by AHA grant 17SFRN33520017. MFL is supported by PO1 HL116263 and R01 HL127173. SS-R is supported by the Frontiers of Innovation Scholars Program (FISP; #3-P3029), the Interdisciplinary Research Fellowship in NeuroAIDS (IRFN; MH081482), and a pilot award from DA037844. SS-R and AAP are supported by funds from the California Tobacco-Related Disease Research Program (TRDRP; Grant Number 28IR-0070). GC is supported by NIH UL1TR000427. JDM is supported by AHA grant 16FTF30130005. Support for NJC was provided by R01 MH113362, U54MD010722, and U01HG009086. This project was conducted in part using the resources of the Advanced Computing Center for Research and Education at Vanderbilt University, Nashville, TN. The datasets used for this project were obtained from Vanderbilt University Medical Center’s Synthetic Derivative, which is supported by numerous sources: institutional funding, private agencies, and federal grants. These include the NIH funded Shared Instrumentation Grant S10RR025141; and CTSA grants UL1TR002243, UL1TR000445, and UL1RR024975.

Data on coronary artery disease / myocardial infarction have been contributed by CARDIoGRAMplusC4D investigators and have been downloaded from www.CARDIOGRAMPLUSC4D.ORG.

The authors thank the staff and participants of the ARIC study for their contributions. ARIC is supported by NHLBI contracts (HHSN268201100005C, HHSN268201100006C, HHSN268201100007C, HHSN268201100008C, HHSN268201100009C, HHSN268201100010C, HHSN268201100011C, and HHSN268201100012C). ARIC/GENEVA was supported by NHGRI grant U01HG004402 (E. Boerwinkle).

### Conflict of Interest Disclosures

PF and SLE are employees of 23andMe, Inc. No other disclosures are reported.

## References

1. Kessler RC, Bromet EJ. The epidemiology of depression across cultures. Annu Rev Public Health. 2013;34:119–138.

2. Walker ER, McGee RE, Druss BG. Mortality in mental disorders and global disease burden implications: a systematic review and meta-analysis. JAMA Psychiatry. 2015;72(4):334–341.

3. Correll CU, Solmi M, Veronese N, et al. Prevalence, incidence and mortality from cardiovascular disease in patients with pooled and specific severe mental illness: a large-scale meta-analysis of 3,211,768 patients and 113,383,368 controls. World Psychiatry. 2017;16(2):163–180.

4. Elovainio M, Hakulinen C, Pulkki-Raback L, et al. Contribution of risk factors to excess mortality in isolated and lonely individuals: an analysis of data from the UK Biobank cohort study. Lancet Public Health. 2017;2(6):e260–e266.

5. Holt-Lunstad J, Smith TB, Baker M, Harris T, Stephenson D. Loneliness and social isolation as risk factors for mortality: a meta-analytic review. Perspect Psychol Sci. 2015;10(2):227–237.

6. Leigh-Hunt N, Bagguley D, Bash K, et al. An overview of systematic reviews on the public health consequences of social isolation and loneliness. Public Health. 2017;152:157–171.

7. Qualter P, Vanhalst J, Harris R, et al. Loneliness across the life span. Perspect Psychol Sci. 2015;10(2):250–264.

8. Matthews T, Danese A, Wertz J, et al. Social isolation, loneliness and depression in young adulthood: a behavioural genetic analysis. Soc Psychiatry Psychiatr Epidemiol. 2016;51(3):339–348.

9. Cacioppo JT, Hawkley LC, Thisted RA. Perceived social isolation makes me sad: 5-year cross-lagged analyses of loneliness and depressive symptomatology in the Chicago Health, Aging, and Social Relations Study. Psychol Aging. 2010;25(2):453–463.

10. Hakulinen C, Pulkki-Raback L, Virtanen M, Jokela M, Kivimaki M, Elovainio M. Social isolation and loneliness as risk factors for myocardial infarction, stroke and mortality: UK Biobank cohort study of 479 054 men and women. Heart. 2018.

11. Kendler KS, Gardner CO, Fiske A, Gatz M. Major depression and coronary artery disease in the Swedish twin registry: phenotypic, genetic, and environmental sources of comorbidity. Arch Gen Psychiatry. 2009;66(8):857–863.

12. Rico-Uribe LA, Caballero FF, Martin-Maria N, Cabello M, Ayuso-Mateos JL, Miret M. Association of loneliness with all-cause mortality: A meta-analysis. PLoS One. 2018;13(1):e0190033.

13. Cuijpers P, Vogelzangs N, Twisk J, Kleiboer A, Li J, Penninx BW. Is excess mortality higher in depressed men than in depressed women? A meta-analytic comparison. J Affect Disord. 2014;161:47–54.

14. Roger VL, Go AS, Lloyd-Jones DM, et al. Heart disease and stroke statistics–2012 update: a report from the American Heart Association. Circulation. 2012;125(1):e2–e220.

15. Xu J, Murphy SL, Kochanek KD, Bastian BA. Deaths: Final Data for 2013. Centers for Disease Control and Prevention;2016.

16. Roden DM, Pulley JM, Basford MA, et al. Development of a large-scale de-identified DNA biobank to enable personalized medicine. Clin Pharmacol Ther. 2008;84(3):362–369.

17. The Atherosclerosis Risk in Communities (ARIC) Study: design and objectives. The ARIC investigators. Am J Epidemiol. 1989;129(4):687–702.

18. Knopman DS, Gottesman RF, Sharrett AR, et al. Mild Cognitive Impairment and Dementia Prevalence: The Atherosclerosis Risk in Communities Neurocognitive Study (ARIC-NCS). Alzheimers Dement (Amst). 2016;2:1–11.

19. Ruderfer DM, Walsh CG, Aguirre MW, Ribeiro JD, Franklin JC, Rivas MA. Significant shared heritability underlies suicide attempt and clinically predicted probability of attempting suicide. Molecular Psychiatry. in press.

20. Delaneau O, Marchini J, Genomes Project C, Genomes Project C. Integrating sequence and array data to create an improved 1000 Genomes Project haplotype reference panel. Nat Commun. 2014;5:3934.

21. Howie BN, Donnelly P, Marchini J. A flexible and accurate genotype imputation method for the next generation of genome-wide association studies. PLoS Genet. 2009;5(6):e1000529.

22. Purcell S, Neale B, Todd-Brown K, et al. PLINK: a tool set for whole-genome association and population-based linkage analyses. Am J Hum Genet. 2007;81(3):559–575.

23. Mosley JD, Shoemaker MB, Wells QS, et al. Investigating the Genetic Architecture of the PR Interval Using Clinical Phenotypes. Circ Cardiovasc Genet. 2017;10(2).

24. Pritchard JK, Stephens M, Donnelly P. Inference of population structure using multilocus genotype data. Genetics. 2000;155(2):945–959.

25. Genomes Project C, Auton A, Brooks LD, et al. A global reference for human genetic variation. Nature. 2015;526(7571):68–74.

26. Yang J, Lee SH, Goddard ME, Visscher PM. GCTA: a tool for genome-wide complex trait analysis. Am J Hum Genet. 2011;88(1):76–82.

27. Carroll RJ, Bastarache L, Denny JC. R PheWAS: data analysis and plotting tools for phenome-wide association studies in the R environment. Bioinformatics. 2014;30(16):2375–2376.

28. Wei WQ, Teixeira PL, Mo H, Cronin RM, Warner JL, Denny JC. Combining billing codes, clinical notes, and medications from electronic health records provides superior phenotyping performance. J Am Med Inform Assoc. 2016;23(e1):e20–27.

29. Breiman L. Random Forests. Machine Learning. 2001;45(1):5–32.

30. Xu H, Stenner SP, Doan S, Johnson KB, Waitman LR, Denny JC. MedEx: a medication information extraction system for clinical narratives. J Am Med Inform Assoc. 2010;17(1):19–24.

31. Whelton PK, Carey RM, Aronow WS, et al. 2017 ACC/AHA/AAPA/ABC/ACPM/AGS/APhA/ASH/ASPC/NMA/PCNA Guideline for the Prevention, Detection, Evaluation, and Management of High Blood Pressure in Adults: A Report of the American College of Cardiology/American Heart Association Task Force on Clinical Practice Guidelines. J Am Coll Cardiol. 2018;71(19):e127–e248.

32. Euesden J, Lewis CM, O’Reilly PF. PRSice: Polygenic Risk Score software. Bioinformatics. 2015;31(9):1466–1468.

33. Wray NR, Ripke S, Mattheisen M, et al. Genome-wide association analyses identify 44 risk variants and refine the genetic architecture of major depression. Nat Genet. 2018;50(5):668–681.

34. Abdellaoui A, Sanchez-Roige S, Sealock J, et al. Phenome-wide investigation of health outcomes associated with genetic predisposition to loneliness. bioRxiv. 2018.

35. Schizophrenia Working Group of the Psychiatric Genomics C. Biological insights from 108 schizophrenia-associated genetic loci. Nature. 2014;511(7510):421–427.

36. Zhu Z, Zheng Z, Zhang F, et al. Causal associations between risk factors and common diseases inferred from GWAS summary data. Nat Commun. 2018;9(1):224.

37. Wyman L, Crum RM, Celentano D. Depressed mood and cause-specific mortality: a 40- year general community assessment. Ann Epidemiol. 2012;22(9):638–643.

38. Cacioppo JT, Cacioppo S, Cole SW, Capitanio JP, Goossens L, Boomsma DI. Loneliness across phylogeny and a call for comparative studies and animal models. Perspect Psychol Sci. 2015;10(2):202–212.

39. Cacioppo JT, Hawkley LC, Berntson GG, et al. Do lonely days invade the nights? Potential social modulation of sleep efficiency. Psychol Sci. 2002;13(4):384–387.

40. Hawkley LC, Thisted RA, Masi CM, Cacioppo JT. Loneliness predicts increased blood pressure: 5-year cross-lagged analyses in middle-aged and older adults. Psychol Aging. 2010;25(1):132–141.

41. Prince M, Patel V, Saxena S, et al. No health without mental health. Lancet. 2007;370(9590):859–877.

42. Cacioppo S, Grippo AJ, London S, Goossens L, Cacioppo JT. Loneliness: clinical import and interventions. Perspect Psychol Sci. 2015;10(2):238–249.

43. Brown EG, Gallagher S, Creaven AM. Loneliness and acute stress reactivity: A systematic review of psychophysiological studies. Psychophysiology. 2018;55(5):e13031.

44. Duncan L, Hanyang S, Gelaye B, et al. Analysis of polygenic score usage and performance across diverse human populations. bioRxiv. 2018.

45. Martin AR, Kanai M, Kamatani Y, Okada Y, Neale B, Daly M. Hidden ‘risk’ in polygenic scores: clinical use today could exacerbate health disparities. bioRxiv. 2018.

