## supplementary for "Genetic risk for major depressive disorder and loneliness in gender-specific associations with coronary artery disease"

### Supplementary Online Content

eMethods

#### Development of a Machine Learning Classifier for CAD

CAD was defined in BioVU subjects by a random forest machine learning classifier[1] that integrated data from across the EHR: inpatient and outpatient billing codes from the International Classification of Diseases, 9th edition (ICD-9), Current Procedural Terminology (CPT) codes, laboratory values, reports, and clinical documentation. The algorithm was developed in a subset of VUMC patients who were born before 1989 and who had complete demographic information (gender, date of birth, race/ethnicity) listed in their clinical record (eFigure 1). We also removed patients for whom classification would be trivial, i.e., those whose EHR lacked any CAD features that were used for classification. From this group of 330,802 patients, we randomly selected training (N= 773) and testing (N=294) sets, and the de-identified EHR of these patients was manually reviewed and adjudicated to either a "CAD" or "No CAD" group. The remaining 329,835 patients were assigned to the implementation set.

The random forest method iteratively constructed decision trees using an *a priori* set of defined features to segregate "CAD" and "No CAD" patients in the training set. Feature weights were developed in the training set and were then used to construct a continuous predicted probability (score) of having CAD for each individual in the testing set, where the performance of the classifier was evaluated. The algorithm was built using the Python package ScikitLearn v0.18.1.[2] Features for algorithm training included 162 CAD-related ICD-9 codes, CPT codes, laboratory values, text strings, and medications. For each feature, 3 metrics were used: counts (the number of times a feature was in a record), persistence (the proportion of record the feature was present), and durability (the proportion of record from the first mention to the end of a record the feature was present). An optimal score threshold (.662) was identified by five-fold cross validation in the testing set, and was selected to yield a false positive rate <1.25% (eFigure 2). The final model evaluated in the testing set had a positive predictive value of .97, a negative predictive value of .90, a sensitivity of .73, and a specificity of .99. Features that were most strongly predictive of CAD in the final model (features with importances > .001)[1] are shown in eFigure 3. Age at CAD diagnosis was defined in cases by the date at first EHR mention of a CAD-related ICD or CPT code, or a non-negated text mention of coronary artery bypass grafting or stent.

Since the negative predictive value of the algorithm was .90, we applied additional filters to the "No CAD" patients before including them in our control group. First, we only included patients with scores < .20, indicating that < .20 of trees had classified the patient as having CAD. Second, we required "No CAD" patients to have a minimum record length of two years and to be at least 60 years of age at last record so that controls would have had an opportunity to develop CAD. We also augmented the control set with patients who did not require classification by the algorithm (i.e., with no EHR evidence of CAD), and who met the above criteria for record length and age at last record.

We genetically validated our algorithm by comparing the SNP-based heritability ($h^2_{SNP}$) of CAD in our case-control sample to $h^2_{SNP}$ estimates that have been previously reported.[3] We estimated $h^2_{SNP}$ by restricted maximum likelihood models in the GCTA package v1.9,[4,5] and compared our estimate to the $h^2_{SNP}$ of the CARDIoGRAMplusC4D Consortium dataset, estimated by LD score regression and reported in LD Hub (http://ldsc.broadinstitute.org/ldhub/).[3] The $h^2_{SNP}$ of CAD in BioVU was .118 (se=.036) on the observed scale, and .107 (se=.033) on the liability scale (computed using a disease prevalence of 6.2%[6]). In comparison, the $h^2_{SNP}$ of CAD in the CARDIoGRAMplusC4D Consortium dataset was .073 (se=.005) on the observed scale. Although our estimate was slightly higher, the difference was not statistically significant (P=.58).

#### Extraction of Covariate Data from the EHR

A strength of the EHR is the availability of data collected over the lifespan on risk factors for common complex diseases, and we included variables for many of these risk factors in our analyses. BMI is frequently calculated at an office visit, resulting in multiple measurements for each patient, which we summarized as a single median value. Hypertension was defined by the presence of ICD-9 codes (401*-405*) and ICD-10 codes (I10*-I13*, I16*), and by problem list mentions of “hypertension” or “htn” (excluding “portal hypertension”, “pulmonary hypertension”, “rv hypertension”, “intracranial hypertension”, and “phtn”). Patients with type 2 diabetes were identified using a BioVU algorithm (https://phekb.org/phenotype/type-2-diabetes-demonstration-project )[7] in which cases had at least one ICD-9 or ICD-10 type 2 diabetes code and mention of non-insulin hypoglycemic medication. Measurements of HDL, LDL, and triglycerides were extracted from laboratory values in the EHR. LDL was calculated by the Friedewald formula.[8] Observations were filtered if they were outside of 1-200 mg/dL for HDL, 1-400 mg/dL for LDL, and 1-1000 mg/dL for triglycerides, which removed less than 4% of observations for each lipid trait. We also excluded observations in patients <18 years of age and any observations on or after the first mention of lipid-lowering medication (eTable 1) in the EHR. Medication status was abstracted from free text in clinical notes via an in-house natural language processing tool.[9] Median lipid values were calculated in patients with multiple observations.

Smoking was defined by data collected on clinic and nursing intake forms, augmented by ICD-9 codes that mapped to the phecode, Tobacco use disorder (318, including all child phecodes; see Phecode Map v1.2: https://phewascatalog.org/phecodes). Individuals identified as “never smokers” were required to have been coded as such on their intake forms, *and* could have no ICD-9 codes indicative of tobacco use disorder. Current or previous smokers were defined as those who had any evidence of smoking in their EHR including clinical notes or ICD-9 codes indicative of tobacco use disorder. Patients for whom a determination of “never smoker” or “current or previous smoker” could not be made were assigned an “unknown” smoking status so that they could be included in multivariable models. Socio-economic status was determined by proxy based on the highest level of education abstracted from the EHR by a natural language processing algorithm.[10] Patients for whom socio-economic status could not be determined by the algorithm were assigned a value of “unknown” so that they could be included in multivariable models.

Psychiatric symptoms were defined by the presence of one or more ICD-9 codes that mapped to phecodes indicative of psychological disorders (295-306.99). Parent phecodes and their descriptions are: Schizophrenia and other psychotic disorders (295), Mood disorders (296), Suicidal ideation or attempt (297), Anxiety disorders (300), Personality disorders (301), Sexual and gender dysphoria (302), Psychogenic and somatoform disorders (303), Adjustment reaction (304), Eating disorder (305.2), and Other mental disorders (306).

#### Development of Polygenic Scores

We calculated polygenic scores via the pruning and thresholding method implemented in PRSice v2.[11] SNPs in the meta-GWAS were pruned using an algorithm that scanned over a 250 kb window and removed SNPs in linkage disequilibrium ($r^2$>0.1) with the most associated SNP (i.e., lowest P), thus retaining the strongest trait-associated SNP within each linkage disequilibrium block. The algorithm cycled through all SNPs in the meta-GWAS, beginning with the SNP with the lowest P and ending at the SNP whose P exceeded a provided threshold, only allowing each SNP to appear in one clump. Linkage disequilibrium estimates were derived directly from the BioVU genotype data, and only pruned SNPs were used in calculating polygenic scores.

In addition to polygenic scores for MDD and loneliness, we constructed a polygenic score for CAD to be used as a covariate in multivariable models. Meta-GWAS summary statistics were obtained from the CARDIoGRAMplusC4D Consortium GWAMA of CAD,[12] and the polygenic score was computed at a P threshold of $5.0005\times10^{-4}$, selected by iterating over *P* thresholds from $5\times10^{-8}$ to 1 in increments of $5\times10^{-5}$ and assessing fit via Nagelkerke's pseudo $R^2$. The polygenic score calculated at this threshold included 684 SNPs, explained 2.11% of the observed phenotypic variance (2.07% on the liability scale), and was strongly associated with CAD risk (OR, 1.46 [95% CI, 1.38-1.55], P=$4.43\times10^{-41}$) per 1-SD increase, in models adjusting for sex, age, batch, and the first 10 principal components of ancestry. Our polygenic score for CAD was calculated at a lower P threshold and included fewer SNPs than recently derived polygenic predictors for CAD.[13,14] Many BioVU patients, however, used lipid-lowering medication, which reportedly offsets a high polygenic risk for CAD,[13] and since the primary purpose of the polygenic score for CAD was to maximize genetic prediction of CAD in our sample, our choice of P threshold was appropriate.

eTable 1. Lipid-lowering medications extracted from the EHR of BioVU subjects using MedEX.[9]

| Medication Class | Brand Name | Generic Name |
|---|---|---|
| bile acid sequestrants | colestid | colestipol |
| bile acid sequestrants | prevalite | colestyramine |
| bile acid sequestrants | questran | colestyramine |
| bile acid sequestrants | welchol | colesevelam |
| cholesterol absorption inhibitors | zetia | ezetimibe |
| fibric acid derivatives | antara | fenofibrate |
| fibric acid derivatives | antara | fenofibrate |
| fibric acid derivatives | clofibrate | fenofibrate |
| fibric acid derivatives | fenoglide | fenofibrate |
| fibric acid derivatives | fibricor | fenofibrate |
| fibric acid derivatives | lipofen | fenofibrate |
| fibric acid derivatives | lofibra | fenofibrate |
| fibric acid derivatives | lopid | gemfibrozil |
| fibric acid derivatives | triglide | fenofibrate |
| fibric acid derivatives | trilipix | fenofibrate |
| nicotinic acid/niacin | niacor | niacin |
| nicotinic acid/niacin | niaspan | niacin |
| PCSK9 inhibitors | praluent | alirocumab |
| PCSK9 inhibitors | repatha | evolocumab |
| probucol | probucol | probucol |
| statin | advicor | niacin extended-release/simvastatin |
| statin | altoprev | extended-release/lovastatin |
| statin | baycol | cerivastatin |
| statin | caduet | amlodipine/atorvastatin |
| statin | canef | fluvastatin |
| statin | crestor | rosuvastatin |
| statin | juvisync | sitagliptin/simvastatin |
| statin | lescol | fluvastatin |
| statin | lescol xl | fluvastatin |
| statin | lipitor | atorvastatin |
| statin | lipobay | cerivastatin |
| statin | liptruzet | ezetimibe/atorvastatin |
| statin | livalo | pitavastatin |
| statin | mevacor | lovastatin |
| statin | pravachol | pravastatin |
| statin | selektine | pravastatin |
| statin | simcor | niacin extended-release/simvastatin |
| statin | statins | statins |
| statin | vastin | fluvastatin |
| statin | vytorin | ezetimibe/simvastatin |
| statin | zocor | simvastatin |

eTable 2. Covariates in the ARIC replication analysis. A description of variables can be found at: ftp://ftp.ncbi.nlm.nih.gov/dbgap/studies/phs000280/phs000280.v3.p1/pheno_variable_summaries/phs000280.v3.pht000114.v2.GENEVA_ARIC_Subject_Phenotypes.data_dict.xml.

| Variable | Description | Units | Coded values |
|---|---|---|---|
| anta07a | Waist girth to nearest cm at visit 1 | cm | |
| bmi01 | Body mass index in kg/(m*m) at visit 1 | kg/(m*m) | |
| cholmdcode01 | Cholesterol-lowering medication within 2 weeks | | 1=Yes<br>0=No |
| cholmdcode02 | Medications which secondarily affect cholesterol | | 1=Yes<br>0=No |
| cigt01 | Cigarette smoking status at visit 1 | | 1=Current smoker<br>2=Former smoker<br>3=Never smoker<br>4=Unknown, but one of the other 3 categories may be ruled out |
| diabts03 | Diabetes with fasting glucose cut point 126mg/dL at visit 1 | | 1=Fasting glucose >=126 mg/dL or non-fasting glucose >=200mg/dL or self-report of physician diagnosis or took diabetic medication in previous two weeks<br>0=All negative |
| elevel01 | Education level, definition 1, at visit 1 | | 1=Grade school or 0 years education<br>2=High school, but no degree<br>3=High school graduate<br>4=Vocational school<br>5=College<br>6=Graduate school or Professional school |
| hdlsiu02 | Re-calibrated HDL cholesterol in mmol/L | mmol/L | |
| hyptmdcode01 | Hypertension lowering medication use; definition 1, at visit 1 | | 1=Yes<br>0=No |
| ldlsiu02 | Re-calibrated LDL cholesterol in mmol/L | mmol/L | |
| sbpa21 | Systolic blood pressure (average of 2nd and 3rd readings) at visit 1 | mm Hg | |
| sbpa22 | Diastolic blood pressure (average of 2nd and 3rd readings) at visit 1 | mm Hg | |
| trgsiu01 | Total triglycerides in mmol/L | mmol/L | |

eTable 3. Characteristics of BioVU CAD cases and controls.

| | CAD cases (N=3893) | CAD controls (N=4197) | *P*[a] |
|---|---|---|---|
| Age, mean (SD) | 63.4 (11.0) | 71.1 (8.0) | <2.20E-16 |
| Female, No. (%) | 1142 (29.3) | 2602 (62.0) | <2.20E-16 |
| MDD diagnosis, No. (%) | 149 (3.8) | 160 (3.8) | 1.96E-01 |
| Depression diagnosis, No. (%) | 958 (24.6) | 1050 (25.0) | 7.49E-01 |
| Other psychiatric disorder diagnosis, No. (%) | 495 (12.7) | 596 (14.2) | 2.77E-02 |
| BMI, mean (SD) | 29.31 (5.73) | 28.71 (6.10) | 2.78E-05 |
| BMI group | | | |
| Normal (BMI<25), No. (%) | 828 (21.3) | 1158 (27.6) | |
| Overweight (BMI≥25 and BMI<30), No. (%) | 1332 (34.2) | 1397 (33.3) | 1.48E-02 |
| Obese (BMI≥30), No. (%) | 1441 (34.2) | 1399 (33.3) | |
| Unknown, No. (%) | 292 (7.5) | 243 (5.8) | |
| Smoking status | | | |
| Never, No. (%) | 1084 (27.8) | 1945 (46.3) | <2.20E-16 |
| Current or former, No. (%) | 1644 (42.2) | 1075 (25.6) | |
| Unknown, No. (%) | 1165 (29.9) | 1177 (28.0) | |
| Hypertension diagnosis, No. (%) | 3715 (95.4) | 3133 (74.6) | <2.20E-16 |
| Type 2 diabetes diagnosis, No. (%) | 426 (10.9) | 238 (5.7) | <2.20E-16 |
| Highest level of education | | | |
| Less than high school, No. (%) | 159 (4.1) | 96 (2.3) | |
| High school, No. (%) | 925 (23.8) | 1108 (26.4) | 5.50E-12 |
| Bachelor's degree, No. (%) | 194 (5.0) | 260 (6.2) | |
| Graduate school, No. (%) | 481 (12.4) | 584 (13.9) | |
| Unknown, No. (%) | 2134 (54.8) | 2149 (51.2) | |
| Antilipemic medication use, No. (%) | 3773 (96.9) | 2479 (59.1) | <2.20E-16 |
| Pre-medication median blood HDL in mg/dL, mean (SD) | 42.7 (14.78) | 56.55 (19.10) | <2.20E-16 |
| Pre-medication median blood LDL in mg/dL, mean (SD) | 117.1 (37.86) | 119.2 (33.08) | 6.87E-01 |
| Pre-medication median triglycerides in mg/dL, mean (SD) | 191.9 (117.23) | 148.1 (99.25) | 7.16E-15 |

[a]*P* from associations in a logistic regression model adjusted for age and sex.

eTable 4. Characteristics of ARIC cases and non-cases. Values were reported at first visit (i.e., "visit 1"), unless otherwise noted.

| | Incident CAD cases (N=923) | Non-cases (N=6274) | $P^a$ |
|---|---|---|---|
| Age at baseline, mean (sd) | 55.6 (5.4) | 53.9 (5.7) | <2.20E-16 |
| Age at event, mean (sd) | 65.1 (6.7) | 69.5 (5.9) | NA |
| Female, N (%) | 287 (31.1) | 3656 (58.3) | <2.20E-16 |
| BMI, mean (sd) | 27.9 (4.70) | 26.7 (4.7) | 1.10E-10 |
| BMI group | | | |
| Normal (BMI<25), N (%) | 239 (25.9) | 2535 (40.4) | |
| Overweight (BMI≥25 and BMI<30), N (%) | 259 (28.1) | 2460 (39.2) | 2.91E-09 |
| Obese (BMI≥30), N (%) | 423 (45.8) | 1275 (20.3) | |
| Unknown | 2 (0.2) | 4 (0.1) | |
| Waist girth in cm, mean (sd) | 100.0 (12.4) | 94.8 (13.2) | 2.26E-13 |
| Smoking status | | | |
| Never, N (%) | 254 (27.5) | 2654 (42.3) | |
| Former, N (%) | 378 (41.0) | 2140 (34.1) | 2.43E-15 |
| Current, N (%) | 291 (31.5) | 2654 (42.3) | |
| Unknown, N (%) | 0 | 4 (0.1) | |
| Hypertension diagnosis[b], N (%) | 507 (54.9) | 2366 (37.7) | <2.20E-16 |
| Systolic blood pressure, mean (sd) | 124.4 (17.9) | 117.2 (16.3) | <2.20E-16 |
| Diastolic blood pressure, mean (sd) | 73.7 (11.0) | 71.4 (9.7) | 1.87E-04 |
| Type 2 diabetes diagnosis, N (%) | 179 (19.4) | 380 (6.1) | <2.20E-16 |
| Highest Level of Education | | | |
| Less than high school | 57 (6.2) | 219 (3.5) | |
| High School | 440 (47.7) | 2777 (44.3) | 4.57E-08 |
| Bachelor's Degree | 353 (38.2) | 2626 (41.9) | |
| Graduate School | 73 (7.9) | 642 (10.2) | |
| Unknown | 0 | 10 (0.2) | |
| Cholesterol-lowering medication use within 2 weeks, N (%) | 41 (4.4) | 176 (2.8) | 1.54E-02 |
| Medications that secondarily affect cholesterol | 240 (26.0) | 1142 (18.2) | 2.94E-09 |
| HDL in mg/dL, mean (sd) | 43.5 (12.9) | 52.6 (17.0) | <2.20E-16 |
| LDL in mg/dL, mean (sd) | 148.7 (36.9) | 134.5 (37.3) | <2.20E-16 |
| Triglycerides in mg/dL, mean (sd) | 163.9 (112.7) | 129.6 (87.6) | <2.20E-16 |

[a]*P* from associations in a Cox proportional hazards model adjusted for baseline age and sex.
[b]Hypertension at baseline in ARIC participants was defined according to recent criteria for the detection of high blood pressure[15] using the criteria: sbpa21>130 or sbpa22>80 or hyptmdcode01=1.

eFigure 1. Development of a random forest (RF) machine learning classifier to identify patients with coronary artery disease, "CAD", and "No CAD" from the Vanderbilt University Medical Center (VUMC) electronic health record (EHR) research database. The machine learning algorithm was trained and tested on manually adjudicated sets of patients with "CAD" and "No CAD". The optimal case selection threshold (.662) was identified by five-fold cross validation in the training set, and performance metrics in the testing set are presented. The final algorithm was deployed in an implementation set to identify CAD cases across the EHR. AUC denotes area under the curve; PPV, positive predictive value.

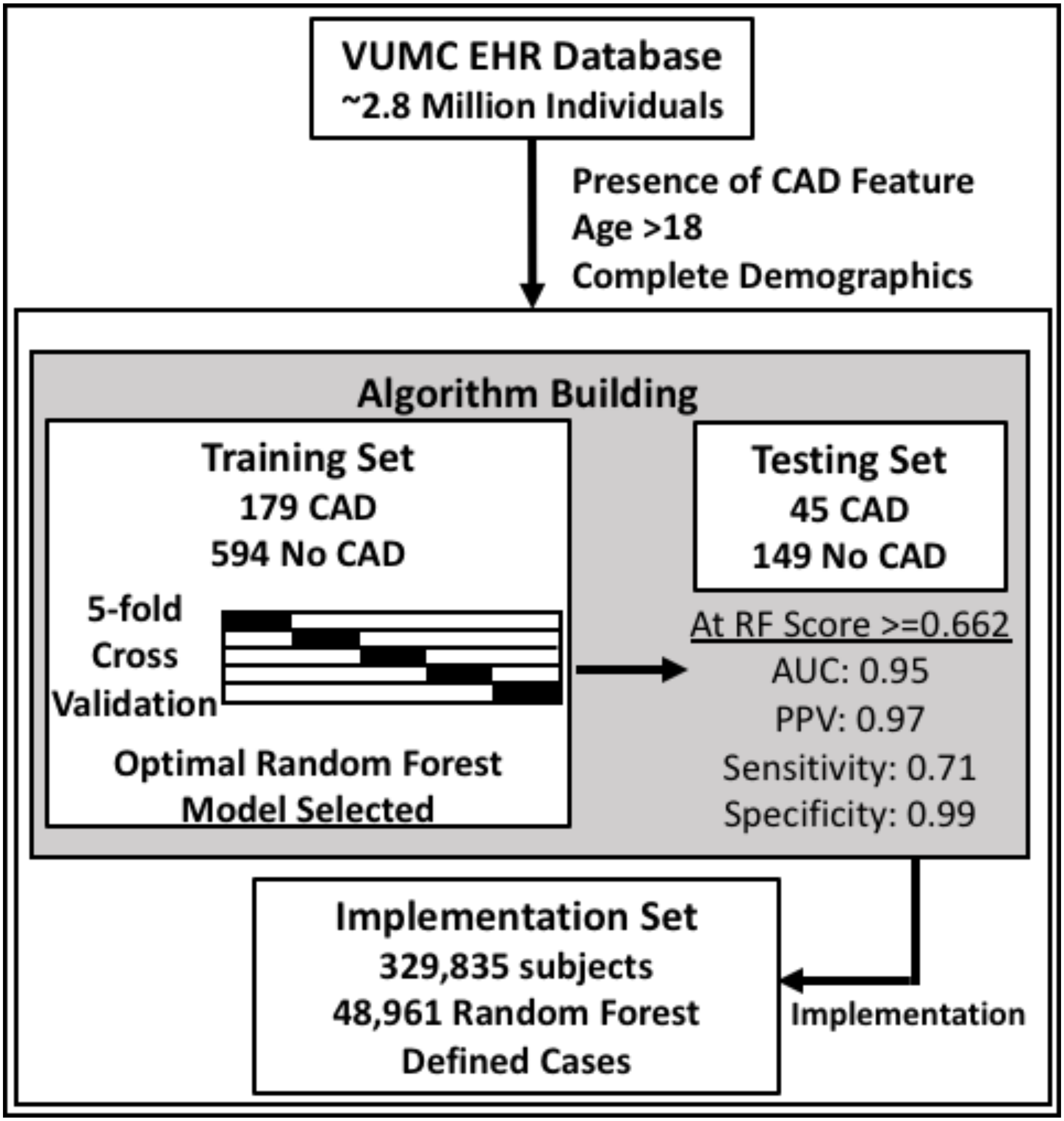

**eFigure 2. Selection of the optimal case score threshold in the random forest machine learning classifier for coronary artery disease.** Five-fold cross validation was used in the training set to optimize tree number and the number of features sampled for splitting at each node. The model with the highest mean out-of-bag area under the curve (AUC) was chosen as the optimal model. Using the mean receiver operating characteristic (ROC) curve in the out-of-bag training set, we chose the minimum score threshold that yielded a false positive rate (FPR) <1.25 % as the case score threshold for the testing and implementation sets.

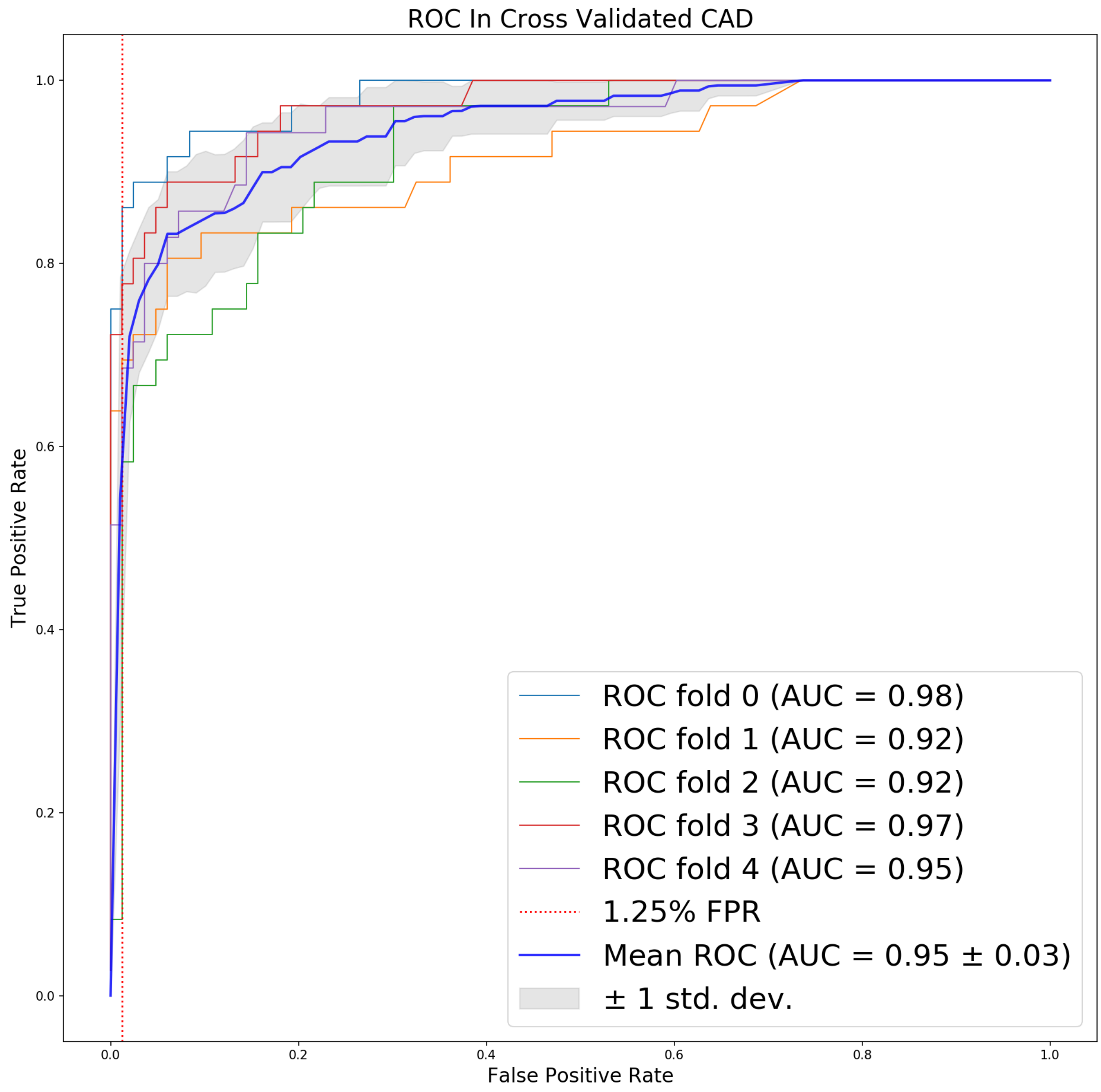

eFigure 3. Importance (weight) of coronary artery disease (CAD) features in the random forest machine learning classifier. Weights were developed in the training set and were used to construct a continuous score for each individual in the testing and implementation sets. Features with a importances > 0.001 are included in the figure.

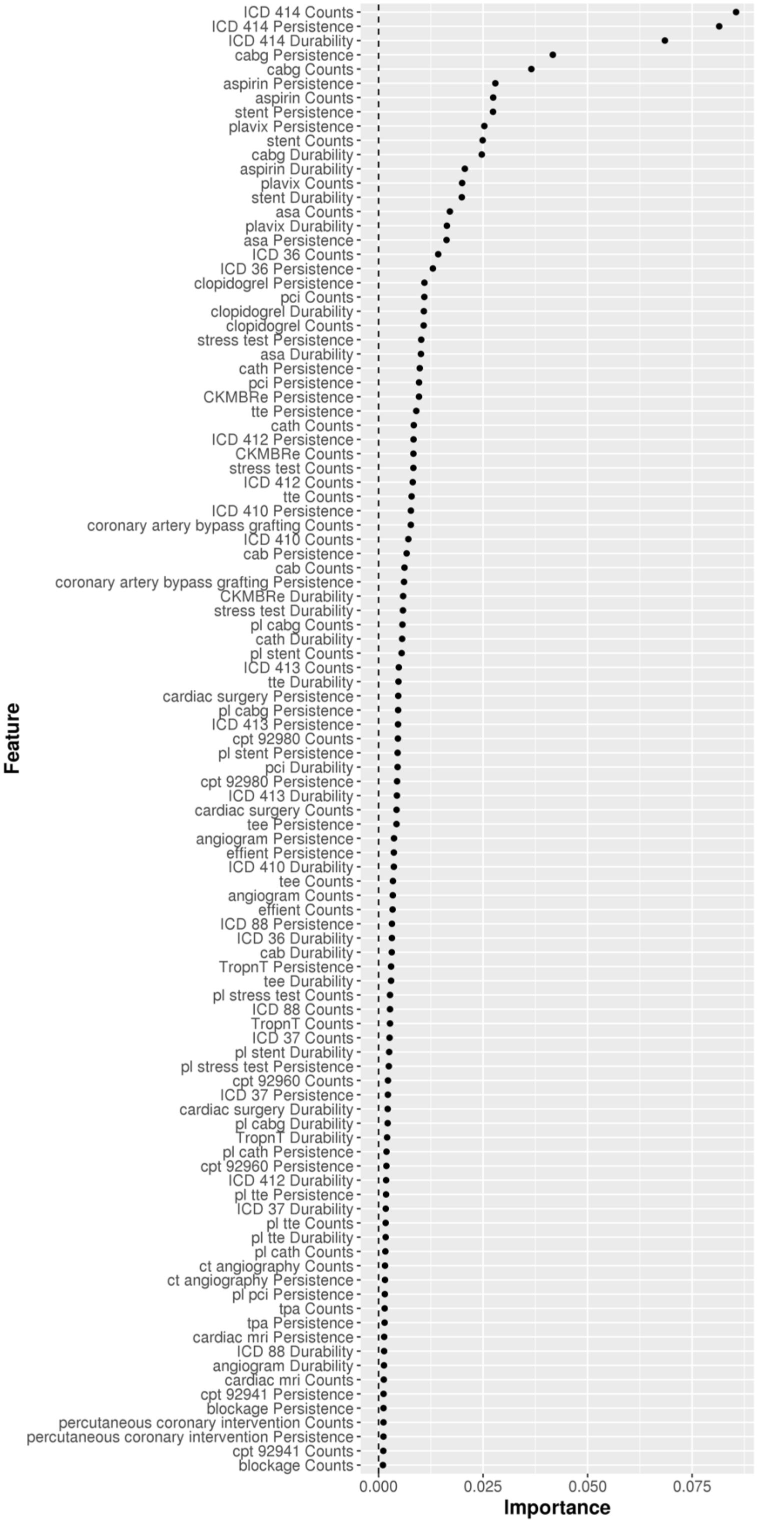

eFigure4. Proportion of variability in coronary artery disease risk explained (Nagelkerke's pseudo $R^2$; observed scale) in BioVU by polygenic scores for major depressive disorder (MDD) and loneliness calculated at P thresholds ranging from $5x10^{-8}$ to .5 in intervals of $5x10^{-5}$. Black dots denote the best-fit P thresholds for each polygenic score.

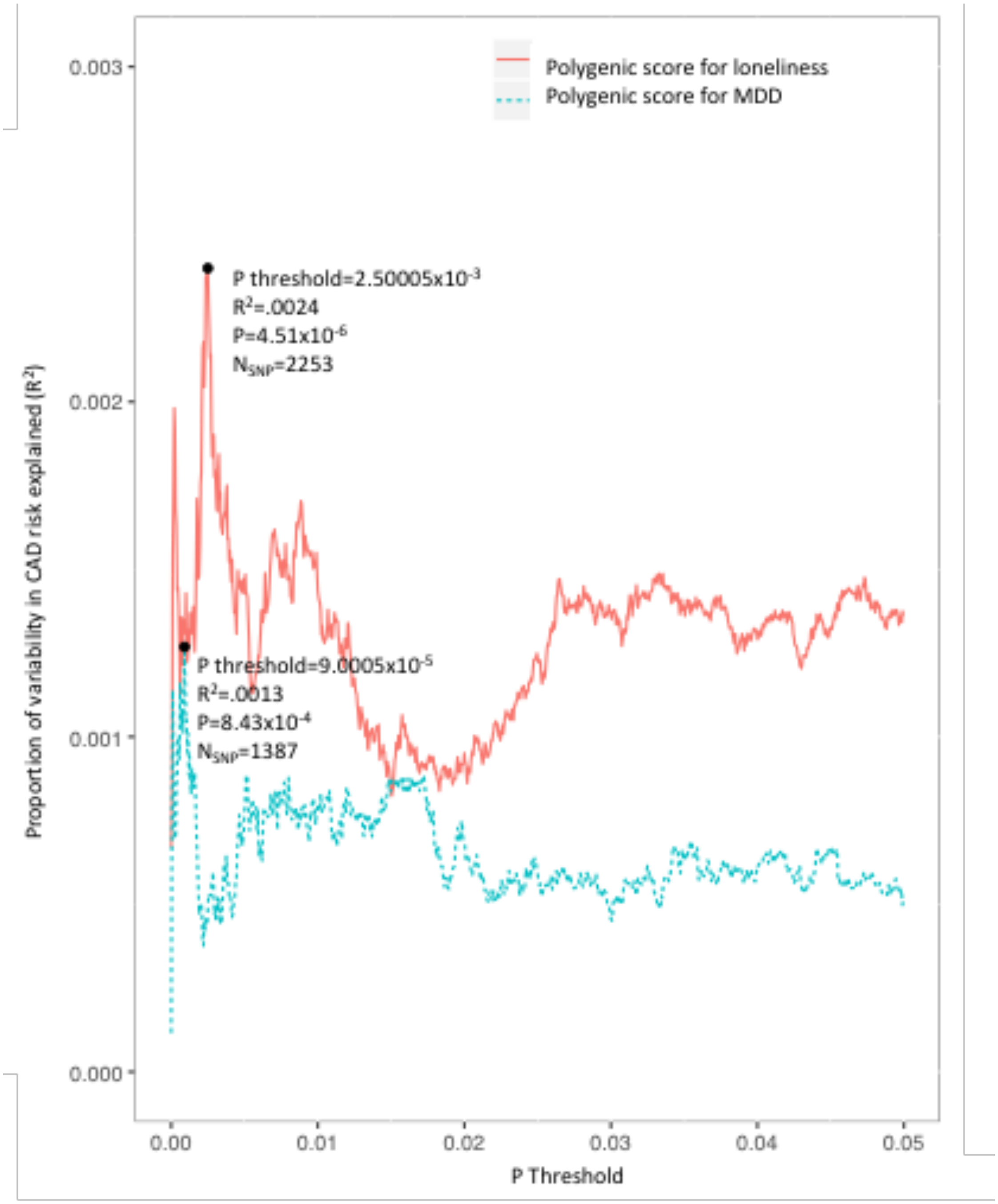

eFigure 5. Risk of CAD in ARIC predicted by polygenic scores for MDD (A), loneliness (B), MDD|loneliness (C), and loneliness|MDD (D) before and after adjustment for conventional cardiovascular disease risk factors. Minimally adjusted ("Minimal") models included sex (except sex-stratified results), age, the first 10 principal components of ancestry, and genotype batch. Fully adjusted ("Full") models included additional covariates for waist girth, smoking status, hypertension medication use, systolic blood pressure, type 2 diabetes, highest level of education, use of cholesterol-lowering medication and other medications that secondarily affect cholesterol, and blood measurements of HDL, LDL, and triglycerides. The asterisk in panel D denotes sex*polygenic score interaction P < .05.

A

| Group | No | No. events | Polygenic score for MDD HR | P |
|---|---|---|---|---|
| All | 7197 | 923 | 1.07 (0.99-1.14) | 0.07 |
| Females | 3943 | 287 | 1.08 (0.95-1.22) | 0.22 |
| Males | 3254 | 636 | 1.06 (0.97-1.15) | 0.19 |
| All | 7009 | 886 | 1.06 (0.99-1.14) | 0.10 |
| Females | 3873 | 275 | 1.09 (0.96-1.25) | 0.17 |
| Males | 3136 | 611 | 1.05 (0.96-1.14) | 0.31 |

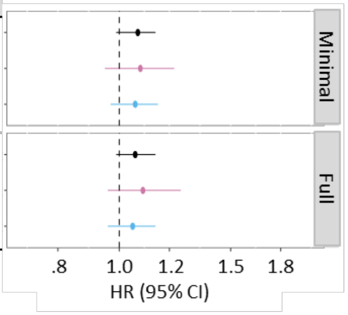

B

| Polygenic score for loneliness HR | P |
|---|---|
| 1.07 (1.01-1.15) | 0.03 |
| 1.17 (1.05-1.32) | 0.01 |
| 1.03 (0.95-1.12) | 0.46 |
| 1.05 (0.99-1.13) | 0.12 |
| 1.12 (1.00-1.26) | 0.05 |
| 1.01 (0.94-1.10) | 0.72 |

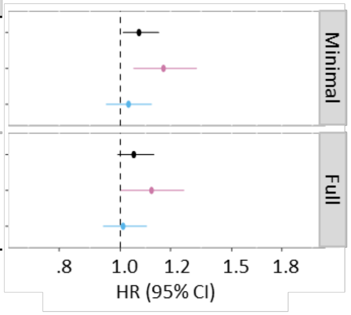

C

| Group | No | No. events | Polygenic score for MDD\|loneliness HR | P |
|---|---|---|---|---|
| All | 7197 | 923 | 1.02 (0.95-1.09) | 0.54 |
| Females | 3943 | 287 | 1.04 (0.93-1.17) | 0.49 |
| Males | 3254 | 636 | 1.01 (0.93-1.10) | 0.79 |
| All | 7009 | 886 | 1.02 (0.95-1.10) | 0.52 |
| Females | 3873 | 275 | 1.08 (0.96-1.22) | 0.19 |
| Males | 3136 | 611 | 1.00 (0.92-1.08) | 0.92 |

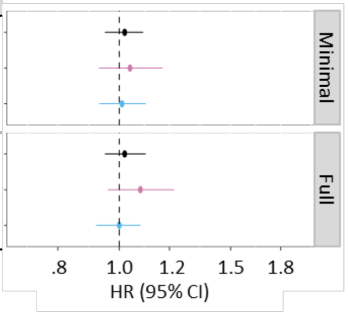

D

| Polygenic score for loneliness\|MDD | |
|---|---|
| 1.04 (0.97-1.11) | 0.25 |
| 1.17 (1.04-1.32) | 0.01 |
| 0.98 (0.91-1.06) | 0.66 |
| 1.03 (0.97-1.10) | 0.34 |
| 1.14 (1.02-1.29) | 0.02 |
| 0.98 (0.90-1.06) | 0.60 |

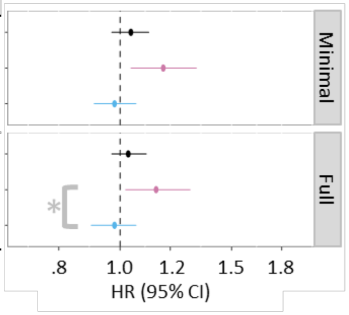

References in Supplementary Material

1. Breiman L. Random Forests. *Machine Learning.* 2001;45(1):5-32.
2. Pedregosa FV, G.; Gramfort, A.; Michel, V.; Thirion, B.; Grisel, O.; Blondel, M.; Prettenhofer, P.; Weiss, R.; Dubourg, V.; Vanderplas, J.; Passos, A.; Cournapeau, D.; Brucher, M.; Perrot, M.; Duchesnay, E.;. Scikit-learn: Machine Learning in Python. *Journal of Machine Learning Research.* 2011;12:6.
3. Zheng J, Erzurumluoglu AM, Elsworth BL, et al. LD Hub: a centralized database and web interface to perform LD score regression that maximizes the potential of summary level GWAS data for SNP heritability and genetic correlation analysis. *Bioinformatics.* 2017;33(2):272-279.
4. Yang J, Lee SH, Goddard ME, Visscher PM. GCTA: a tool for genome-wide complex trait analysis. *Am J Hum Genet.* 2011;88(1):76-82.
5. Lee SH, Wray NR, Goddard ME, Visscher PM. Estimating missing heritability for disease from genome-wide association studies. *Am J Hum Genet.* 2011;88(3):294-305.
6. Writing Group M, Mozaffarian D, Benjamin EJ, et al. Heart Disease and Stroke Statistics-2016 Update: A Report From the American Heart Association. *Circulation.* 2016;133(4):e38-360.
7. Ritchie MD, Denny JC, Crawford DC, et al. Robust replication of genotype-phenotype associations across multiple diseases in an electronic medical record. *Am J Hum Genet.* 2010;86(4):560-572.
8. Friedewald WT, Levy RI, Fredrickson DS. Estimation of the concentration of low-density lipoprotein cholesterol in plasma, without use of the preparative ultracentrifuge. *Clin Chem.* 1972;18(6):499-502.
9. Xu H, Stenner SP, Doan S, Johnson KB, Waitman LR, Denny JC. MedEx: a medication information extraction system for clinical narratives. *J Am Med Inform Assoc.* 2010;17(1):19-24.
10. Hollister BM, Restrepo NA, Farber-Eger E, Crawford DC, Aldrich MC, Non A. Development and Performance of Text-Mining Algorithms to Extract Socioeconomic Status from De-Identified Electronic Health Records. *Pac Symp Biocomput.* 2017;22:230-241.
11. Euesden J, Lewis CM, O'Reilly PF. PRSice: Polygenic Risk Score software. *Bioinformatics.* 2015;31(9):1466-1468.
12. Nikpay M, Goel A, Won HH, et al. A comprehensive 1,000 Genomes-based genome-wide association meta-analysis of coronary artery disease. *Nat Genet.* 2015;47(10):1121-1130.
13. Khera AV, Chaffin M, Aragam KG, et al. Genome-wide polygenic scores for common diseases identify individuals with risk equivalent to monogenic mutations. *Nat Genet.* 2018;50(9):1219-1224.
14. Abraham G, Havulinna AS, Bhalala OG, et al. Genomic prediction of coronary heart disease. *Eur Heart J.* 2016;37(43):3267-3278.
15. Whelton PK, Carey RM, Aronow WS, et al. 2017 ACC/AHA/AAPA/ABC/ACPM/AGS/APhA/ASH/ASPC/NMA/PCNA Guideline for the

Prevention, Detection, Evaluation, and Management of High Blood Pressure in Adults: A Report of the American College of Cardiology/American Heart Association Task Force on Clinical Practice Guidelines. *J Am Coll Cardiol.* 2018;71(19):e127-e248.
